# CUT&TIME captures the history of open chromatin in developing neurons

**DOI:** 10.1101/2025.08.29.673195

**Authors:** Kiara C. Eldred, Matthew Wooten, Derek H. Janssens, Joshua Hahn, Shane J Neph, Sierra J. Edgerton, Gracious Wyatt-Draher, Stephanie M. Sherman, Jane E. Ranchalis, Andrew B. Stergachis, Thomas A. Reh, Steven Henikoff

**Author notes:** Correspondence should be addressed to S.H. and T.A.R. Denotes equal contributions as first author (Kiara C. Eldred, Matthew Wooten).

## Abstract

Chromatin structure plays a central role in defining cell identity by regulating gene expression. During development, shifts in chromatin structure facilitate changes in gene expression needed to specify distinct cell types. To understand how changes in chromatin structure influence the developmental trajectory of neural progenitor cells, we developed CUT&TIME, a technique that uses a hyperactive 6-methyl adenosine (6mA) methyltransferase pulsed in living cells to map historical chromatin accessibility genome-wide in single cells. We show that CUT&TIME produces a record of the chromatin landscape during neurogenesis in the developing retina, specifically as neural progenitors produce the major projection neuron type, retinal ganglion cells (RGCs). We further show that this method is compatible with single cell profiling technologies, which allows us to visualize and capture the diversity of chromatin states that produce RGCs. Additionally, we identify changes in promoter accessibility associated with the transition from progenitor to RGC. Together, these data demonstrate that CUT&TIME captures a historical record of chromatin structure, which can be used to identify early changes in accessibility associated with cell-fate commitment.

## Introduction

An important, unanswered question in developmental biology involves how a cell progresses from a multipotent progenitor to a highly refined cell subtype. Much of our understanding of cell fate specification has been gained by direct observation of lineage tracing during development. Sparse labeling with an inheritable marker gene enables identification of the progeny of individual progenitors, shedding light onto the diversity of differentiated progeny a progenitor is competent to make^1-7^. While powerful, lineage tracing fails to identify genetic elements that drive a progenitor cell to take on a specific cell fate. Pseudotime trajectories built from single cell-compatible methods such as RNA-seq or ATAC-seq have enabled interrogation of genetic elements involved in cell fate decisions, and have yielded valuable insights into the order of gene expression events required to specify many distinct cell types. However, differences in chromatin structure that distinguish self-renewing progenitors from progenitors committed to differentiation remain unclear.

Directed DNA modification has been employed to create a “memory” of open chromatin regions or localization of specific elements in living cells. These methods utilize various bacterial DNA methylation modifications that are not present in the human genome, including m6A^8,9^ and CmeC(A/T)GG penta-nucleotides^10^. While innovative, these methods lack the high-throughput sequencing capability of single cell analysis to get a deeper understanding of the diversity of cell states through developmental time.

To address these challenges, we developed CUT&TIME, or <u>C</u>leavage <u>U</u>nder <u>T</u>argets & <u>T</u>argeted Integration via <u>M</u>ethyltransferase <u>E</u>xpression, a method that interrogates the history of accessible chromatin in differentiated cells at the single cell level. Using a hyperactive nonspecific m6A methyltransferase^11^ (m6A-MTase) expressed in living cells, our method labels open chromatin regions of a progenitor cell with an indelible mark (m6A) that persists throughout the process of cell fate specification. Using a modified sciCUT&TAG protocol, we profile the location of the m6A modification at the single cell level, thereby allowing us to capture the diversity of chromatin states that are competent to produce specific subtypes of differentiated progeny. Using human retinal organoids derived from embryonic stem cells, we demonstrate that CUT&TIME is effective at capturing a record of chromatin changes during human development. By transiently expressing a m6A-MTase to mark accessible chromatin in retinal progenitor cells, CUT&TIME allows us to directly visualize the chromatin landscape that existed in retinal progenitor cells prior to undergoing cell fate specification into neurons. By comparing the historical chromatin structure of RGCs to current RGCs and progenitor cells, CUT&TIME allows for identification of promoters with elevated accessibility in cells that transition from progenitors to RGCs and identification of candidate genes that may be involved in the earliest stages of human RGC specification.

## Results

### m6A-MTase expression deposits m6A in open chromatin regions of live cells

Retinal organoids represent an ideal model to study the molecular mechanism governing cell fate commitment in humans. Stem cell derived human retinal organoids recapitulate human development, and allow access to retinal progenitor cells that are making early cell fate specification decisions^12,13^. During retinal development, a common pool of progenitors produce specific sub-types of retinal cells in dedicated temporal windows, with early-born cell types produced first, and late-born cell types produced later in development^14^. At early time points in retinal development, the retinal progenitors are either proliferating or differentiating into the first-born retinal cell type: RGCs. We focused on this early decision to develop and validate the CUT&TIME method.

To mark the open chromatin of a live cell, we employed the use of a nonspecific hyperactive m6A-methyltransferase (m6A-MTase)^11^. This m6A-MTase was expressed with a lentiviral vector in live retinal organoids derived from stem cells behind a ubiquitous EF1-alpha promotor (**Fig. 1A**). To assess if there was appreciable deposition of the m6A mark on genomic DNA, we performed a dot-blot comparing uninfected organoids, and organoids infected with the m6A-MTase (**Fig. 1B**). We were able to detect m6A deposition with as little as 3.2 ng of genomic DNA derived from infected organoids expressing the m6A-MTase, with little to no detection of m6A deposition in non-transfected controls. Additionally, when we stain nuclei of both infected and non-infected cells for m6A, we see signal specific to a subset of nuclei (**Fig. S1A**).

**Figure 1:**
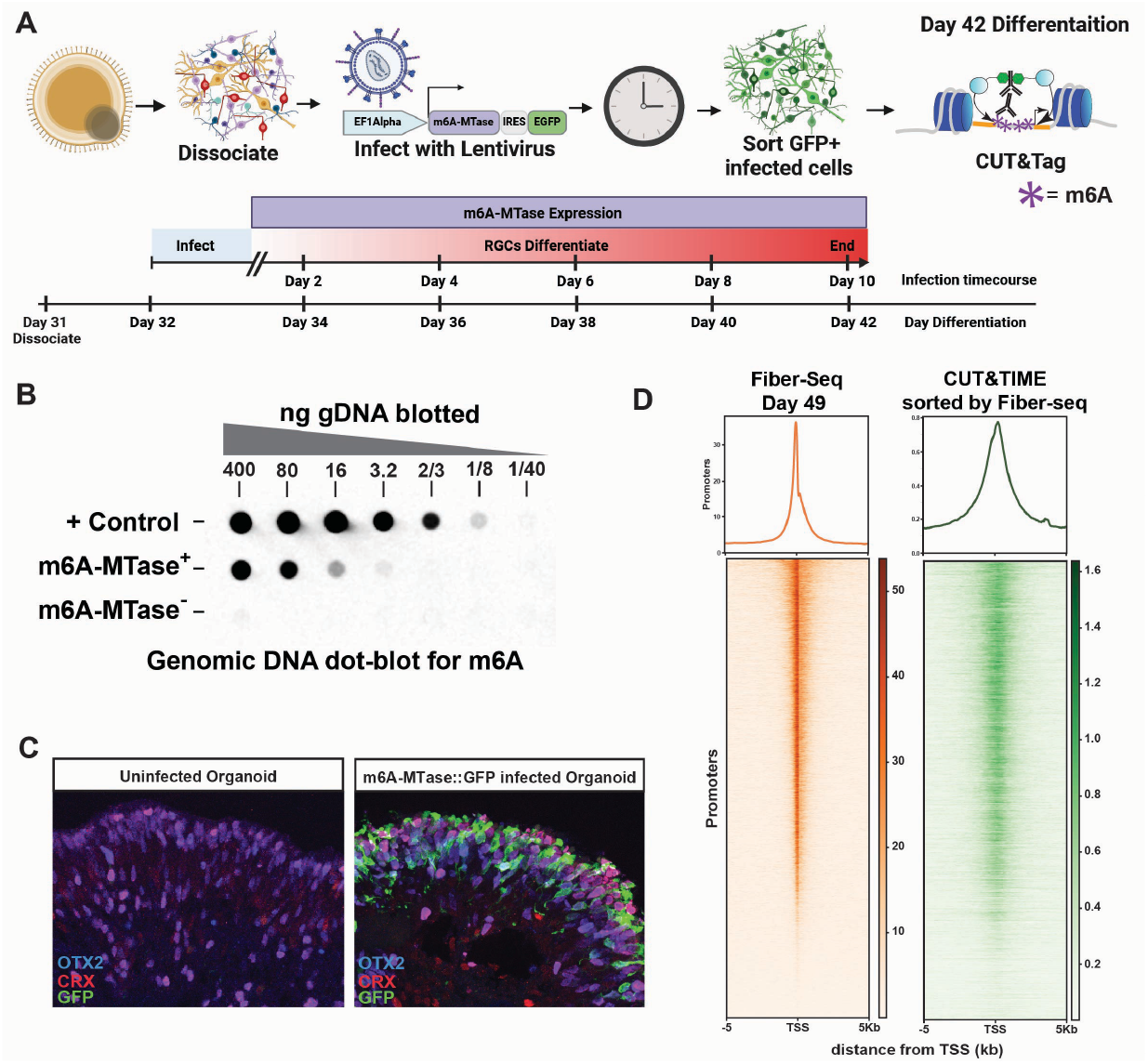
m6A-MTase expression deposits m6A in open chromatin regions of live cells. **(A)** Schematic of experimental set up. **(B)** Dot blot of control bacterial DNA with m6A deposited, genomic DNA extracted from m6A-MTase infected retinal cells, and retinal cells without infection.(C) Day 86 organoids stained for OTX2 (blue), CRX (red), and GFP (green). Left, uninfected organoid. Right, organoid infected at day 26 with m6A-MTase::GFP virus. **(D)** Left: Heat map of fiber-seq signal from plated retinal organoids, treated with a similar experimental protocol as shown in A above, however without m6A infection. 5Kb on either side of all promotors is shown, ordered from most coverage to least coverage. Right: m6A profiling from CUT&TIME protocol with promotors presented in the same order as those from the fiber-seq. Graphs at the top display the average distribution of the m6A modification along the promotors.

The human genome does not endogenously deposit m6A on DNA, however it is possible that the presence of this mark could change the behavior of cells or their gene expression profiles. To test the impact of m6A deposition on retinal development, we expressed m6A-MTase in organoids for 60 days during development and performed immunofluorescent labeling to visualize known markers of cell type specification. We stained organoid sections for OTX2 (bipolar and photoreceptor marker), CRX (bipolar and photoreceptor marker) and GFP (marking lentiviral infection and m6A-MTase expression) (**Fig. 1C**). Both infected and uninfected organoids contained a well differentiated photoreceptor layer, with both infected and uninfected cells showing clear markers of differentiation. We also observed that hESCs expressing the m6A-MTase under the ubiquitous EF1-alpha promoter exhibited normal stem cell morphology and maintained their proliferative potential in the presence of the m6A mark (**Fig. S1B**). Lastly, we profiled the distribution of RNA Polymerase II (Pol II) on the DNA with CUT&Tag, and found no difference in the occupancy of transcription start sites (TSSs) between cells that contained m6A-MTase, and those that did not (**Fig. S1C**). Based on these observations, we conclude that retinal cells can differentiate properly, progenitors can continue to proliferate, and RNA Pol II distribution is normal in cells containing the m6A modification on genomic DNA.

To profile the location of the m6A marks on the genome, we adapted the CUT&Tag protocol to enable detection of the m6A modification. Under normal CUT&Tag conditions, there is no appreciable detection of m6A modification. (**Fig. S1D**). This is because the epitope recognized by the m6A antibody is masked in the presence of double-stranded DNA, meaning the antibody will only bind m6A in regions containing single-stranded DNA. To circumvent this issue, we heat-treated cells for 20 minutes at 98°C to melt the double helix and expose the m6A epitope. We then slowly cooled the samples while adding the primary m6A antibody, similar to methods described for BrdU labeling^15^. This regime permits the antibody to bind to its target while also allowing the melted DNA to reanneal, making it a suitable substrate for Tn5 integration according to the standard CUT&Tag protocol. This augmented protocol called CUT&TIME yields a substantial increase in tagmentation, allowing for effective detection of the m6A modification (**Fig. S1D**).

If the m6A-MTase is behaving similarly in live cells compared to activity in permeabilized nuclei before DNA extraction^11^, we would expect to see deposition of m6A in open chromatin regions, corresponding to those observed with traditional Fiber-seq methods^11^ or with ATAC-seq in similar organoid samples. When m6A profiles are compared directly to those generated by Fiber-seq within 5kb of the transcription start site, we indeed see a similar distribution of signal in both data sets (**Fig. 1D**). When comparing m6A profiles with ATAC-seq from day 49 organoid samples^16^, we again see a similar distribution of signal in these data (**Fig. S1E**).

From these results, we conclude that the m6A-MTase can be expressed in live cells to report on the open chromatin regions, and neither the presence of the enzyme nor the existence of the m6A modification interferes with normal progenitor proliferation or differentiation

### Single cell CUT&TIME resolves distinct cell types in human retinal organoids

To determine if the m6A profile established in live cells can be used to distinguish unique cell types, we adapted CUT&TIME to gain single cell resolution (scCUT&TIME). Retinal organoids were grown to day 35 of differentiation, dissociated, and then infected with a lentivirus containing the m6A-MTase behind an inducible TREG promotor. Cells were then allowed to further differentiate for 10 days, after which doxycycline was added to the medium for two days to drive transcription of the m6A-MTase, resulting in m6A deposition for the final two days of the experiment (**Fig. 2A**). We refer to this labeling regimen as the “late pulse” and this protocol of labeling should provide a snapshot of the accessible chromatin in all the cells in the organoid, much like Fiber-seq or ATAC-seq.

**Figure 2:**
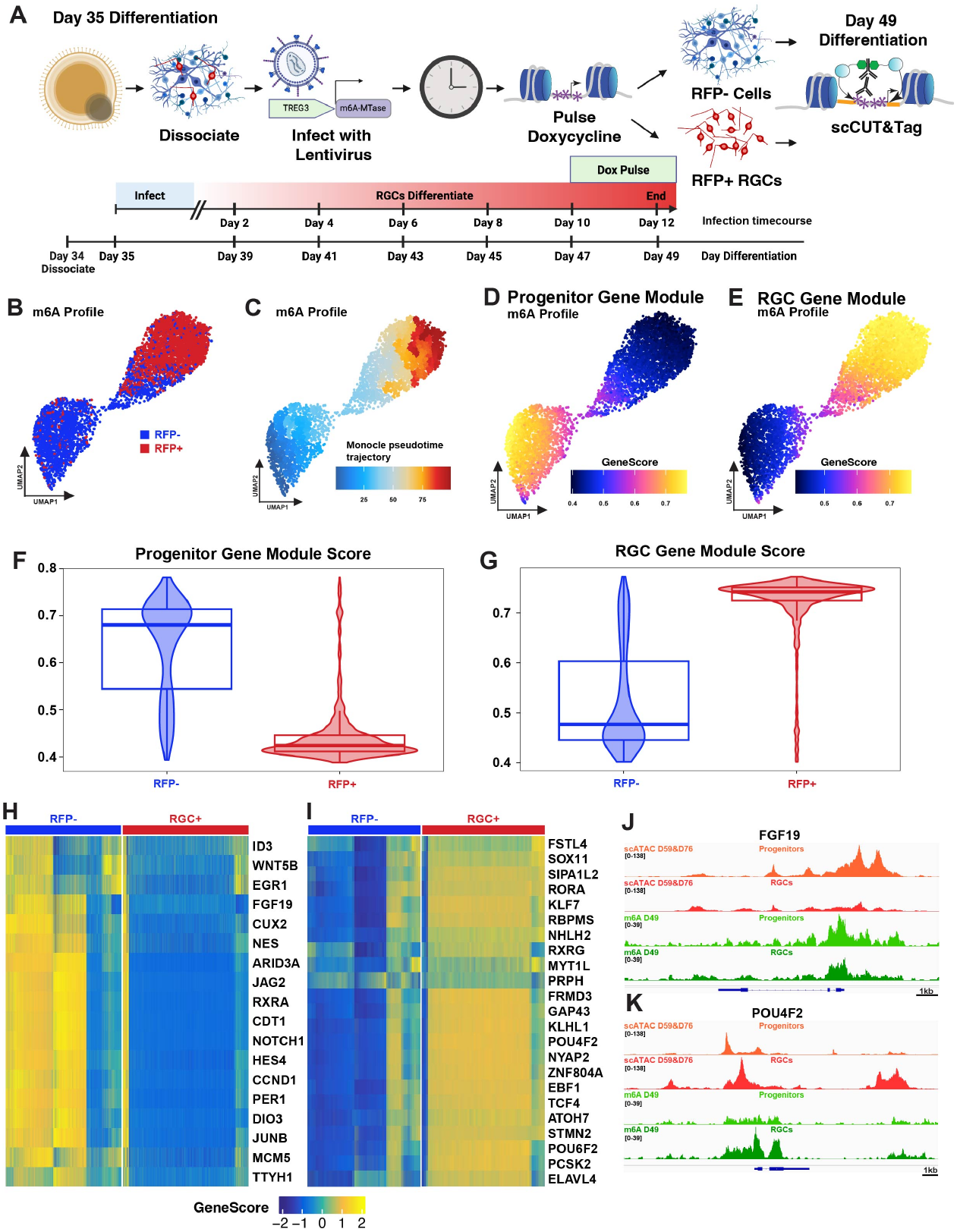
Single cell CUT&TIME distinguishes cell types. **(A)** Schematic of experimental design. **(B)** UMAP of m6A signal generated by CUT&TIME. Cells were sorted based on RFP signal into two groups: RFP+ ganglion cells (red), and RFP-cells (blue). **(C)** Pseudotime using Monocle displaying the most likely path of differentiation as cells progress from progenitors to RGCs. **(D)** Progenitor Gene Module displaying the average m6A signal at genes found in progenitors, listed below in **Fig. 2H. (E)** RGC Gene Module displaying the average m6A signal at genes found in RGCs, listed below in **Fig. 2I.** (F) Violin plot displaying the average progenitor gene module score for each sample. **(G)** Violin plot displaying the average RGC gene module score for each sample. **(H-I)** Heatmaps of m6A signal for genes, displayed for all cells in both the RFP-(blue, left) and RFP+ (Red, right). **(H)** Progenitor genes. (I) RGC genes. **(J-K)** Browser tracks of genomic accessibility as defined by scATAC-seq of fetal retina in progenitors (orange) and RGCs (red) compared to accessibility as profiled by CUT&TIME in progenitors (light green) and RGCs (dark green). **(J)** FGF19, a gene expressed in progenitors. **(K)** POU4F2, a gene expressed in RGCs.

Organoid cultures were then dissociated and sorted using FACS based on POU4F2-driven RFP signal to isolate the RGCs. Cells were then subjected to scCUT&TIME, involving application of a unique barcode to each cell through combinatorial indexing. These samples were then merged in ArchR and a UMAP dimensional reduction was obtained. In the merged UMAP, we see distinct separation between the two sorted cell groups of RFP+ RGCs (red) verses RFP-cells (blue) (**Fig. 2B**). The RFP-cells at this time point should be mainly comprised of retinal progenitor cells. These data demonstrate that CUT&TIME is an effective method to resolve cell types according to their distinct m6A profiles.

To further explore the CUT&TIME technique, we used Monocle to order the cells by pseudotime. Monocle pseudotime trajectory calculates the probability of cells transitioning from one state to another and aligns the cells within transition states according to their gene expression^17^. If we map a Monocle pseudotime trajectory onto these cells using the bottom left of the RFP-cells as the root, we see that the cells mature as they progress towards the distal tip of the RFP+ RGC cluster, as would be expected (**Fig. 2C**). To verify the cell identity of our distinct clusters, we visualized the average ArchR-generated gene scores of known retinal progenitor cells and RGC markers. When we assay genes known to be active in progenitors, we see that if we average the reads within a set of known progenitor genes displayed as a gene score module, the RFP-progenitor cluster is clearly more accessible in these regions compared to the RFP+ RGCs (**Fig. 2D**,**F**, individual gene scores for each gene displayed in **Fig. 2H**). Elevated accessibility at progenitor markers can also be visualized in browser tracks comparing the CUT&TIME signal for RFP-progenitors and RFP+ RGCs across the FGF19 locus, similar to differences in accessibility captured by scATAC data sets previously generated from fetal day 59 and 76^18^ (**Fig. 2J**). Conversely, when we sample gene scores for genes known to be expressed in RGCs, we see elevated gene score module accessibility in the RFP+ RGC population compared to the RFP-progenitor population (**Fig. 2E**,**G**, individual gene scores for each gene displayed in **Fig. 2I**). Elevated accessibility at RGC markers can also be visualized in tracks comparing CUT&TIME signal for RFP-progenitors and RFP+ RGCs across the POU4F2 locus, similar to differences in accessibility captured by scATAC data from fetal day 59 and 76^18^ (**Fig. 2K**). Interestingly, though accessibility differences between progenitors and RGCs are similar when assessed by scCUT&TIME or scATAC-seq, the apex of the peaks generated by scATAC-seq vs. scCUT&TIME are not in the same genomic locations, suggesting subtle differences in the detection preference for accessible sites between these two methods.

### CUT&TIME captures the history of progenitor chromatin structure in differentiated RGCs

We next used CUT&TIME to mark the historical open chromatin of a progenitor cell prior to differentiation into an RGC. To do so, we added doxycycline to the medium for two days, early in organoid development to induce expression of the m6A-MTase and deposition of the m6A mark at open chromatin. The doxycycline was then removed and cells were allowed to differentiate for an additional 8 days. We refer to this labeling regime as the “early pulse” (**Fig. 3A**). Cells were then sorted based on POU4F2-driven RFP into RFP+ RGCs or RFP-progenitor cells.

**Figure 3:**
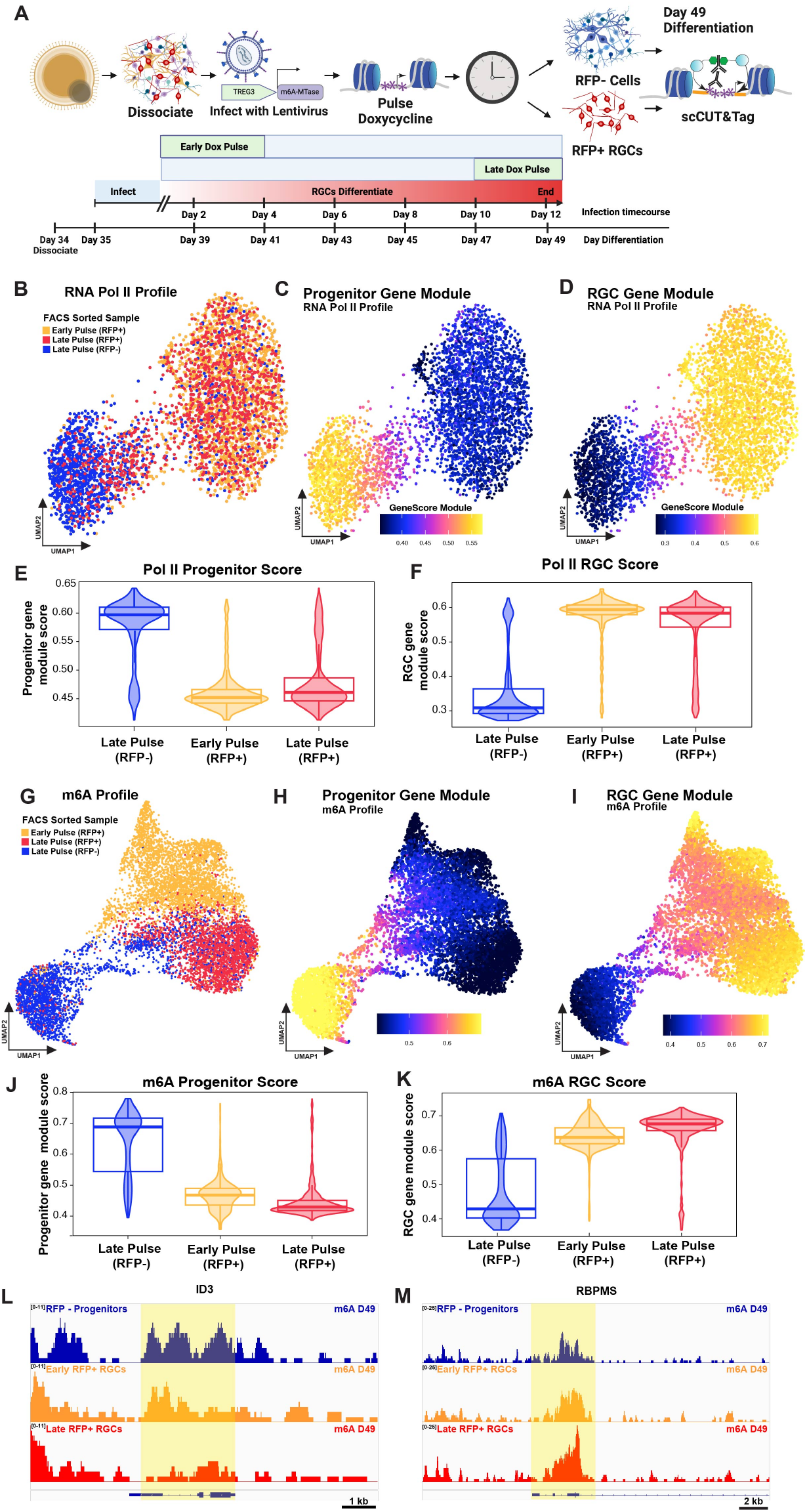
Early pulse of m6A-MTase in progenitor cells leaves a lineage history of open chromatin landscapes during retinal ganglion cell differentiation. **(A)** Schematic of experimental design. **(B-D)** UMAP of RNA Pol II profiles generated by CUT&Tag for retinal cells cultured with experimental protocol displayed in **Fig. 3A. (B)** Three different FACS sorted populations displayed: Early pulse RFP+ ganglion cells (orange), Late pulse RFP+ ganglion cells (red), and Late pulse RFP-progenitor cells (bule). **(C)** Progenitor gene module displaying the RNA Pol II profile at progenitor genes displayed in **Fig. 2F. (D)** RGC gene module displaying the RNA Pol II profile at progenitor genes displayed in **Fig. 2G. (E)** Violin plot summarizing the average progenitor gene module score for RNA Pol II profiles generated by CUT&Tag in each sample shown in **Fig. 3B. (F)** Violin plot summarizing the average RGC gene module score for RNA Pol II profiles generated by CUT&Tag in each sample shown in **Fig. 3B. (G-I)** UMAP of m6A profiles generated by CUT&TIME for retinal cells cultured with experimental protocol displayed in **Fig. 3A. (G)** Three different FACS sorted populations displayed: Early pulse RFP+ ganglion cells (orange), Late pulse RFP+ ganglion cells (red), and Late pulse RFP-progenitor cells (bule). **(H)** Heatmap of m6A signal for progenitor genes, displayed for all cells in both the RFP-(blue, left) and Progenitor gene module displaying the m6A profile at progenitor genes displayed in **Fig. 2H. (I)** RGC gene module displaying the m6A profile at RGC genes displayed in **Fig. 2I. (J)** Violin plot summarizing the average progenitor gene module score for m6A profiles in each sample shown in **Fig. 3G. (K)** Violin plot summarizing the average RGC gene module score for m6A profiles in each sample shown in **Fig. 3G. (L-M)** Browser tracks of genomic accessibility profiles for three cell clusters displayed in **Fig. 3E** showing m6A accessibility. **(L)** displays progenitor gene ID3. **(M)** displays RGC gene RBPMS.

To verify the similarity of chromatin states in FACS sorted RGCs from early pulse and late pulse experiments, we performed sciCUT&Tag targeting Pol II and generated a merged UMAP. When observing the distribution of these samples in UMAP space, we see that Pol II signal is able to clearly separate RFP-progenitors cells from RFP+ RGCs (**Fig. 3B**). As expected, we saw no difference in the clustering of the early vs. late pulsed RFP+ RGCs, indicating that their current chromatin landscapes as marked by Pol II were highly similar. When the progenitor or the RGC gene score modules were applied to this object displaying the averages of genes listed in **Fig. 2 H and I**, we observed higher Pol II levels at the progenitor genes in the RFP-progenitor cluster, and low Pol II levels at progenitor genes in the RFP+ RGCs (**Fig. 3C**). Specific examples for individual gene scores of profiling density along the progenitor genes NOTCH1, DIO3, and ARID3A, are shown in **Fig. S2A**. Conversely, when we applied the RGC gene score Module to this object, displaying an average of the genes listed in **Fig. 2I**, we find that both early and late pulsed RFP+ RGCs have high Pol II binding at RGC genes (**Fig. 3D**). Specific examples for individual gene scores of profiling density along the progenitor genes RBPMS, POU4F2, and EBF1, are shown in **Fig. S2B**. Violin plots show the average gene module scores for the three clusters displayed in **Fig. 3B** (**Fig. 3E-F**). These data indicate that the active gene expression assayed by Pol II profiling in both early and late pulsed RFP+ RGCs is similar, and consistent with RGC fate.

To determine whether we can capture the history of open chromatin as the progenitors generate RGCs, we compared the early and late pulsed RGCs. In principle, if the historical chromatin label is retained as progenitors differentiate into RGCs, the resulting early pulse cohort of RGCs will retain m6A on progenitor genes. We see that unlike the Pol II UMAP described above, the early pulsed RFP+ RGCs, in which the m6A-MTase was active while cells were differentiating from progenitor cells to RGCs, cluster distinctly from the late pulsed RFP+ RGCs (**Fig. 3G**). This differential clustering can be attributed to the higher labeling of progenitor genes in the early pulsed RFP+ RGCs than the late pulsed RFP+ RGCs, as displayed by the progenitor gene module (**Fig. 3H**). Specific examples for individual gene scores of profiling density along the progenitor genes NOTCH1, DIO3, and ARID3A, are shown in (**Fig. S2C**). We next assessed whether the genes in the RGC gene score module show m6A labeling in RGCs of the combined UMAP. In this case, we see that the early pulse RGCs show lower accessibility at RGC genes when compared to the late pulse RGCs (**Fig. 3I**) Specific examples for individual gene scores for the RGC genes RBPMS, POU4F2, and EBF1 are shown in (**Fig. S2D**). Violin plots show the average gene module scores for the three clusters displayed in **Fig. 3G** (**Fig. 3J-K**). These trends can also be seen in individual browser tracks for the progenitor gene ID3 and the RGC gene RBPMS (**Fig. 3L-M**)

To assess the robustness of our method, we performed an additional early pulse experiment in which the cells were collected at day 61 (D61) (**Fig. S2E**), and created a merged UMAP with the late pulse RGCs taken at D49 (**Fig. S2F**). Similar to previous results, we see that the D61 early pulse RGCs cluster distinctly from the late pulse RGCs in the UMAP build from m6A profiling (**Fig. S2F**). When we apply the progenitor and RGC gene score modules to this UMAP, we again see that early pulse D61 RGCs show higher signal for progenitor genes and lower signal for RGC genes when compared to the late pulse RGCs (**Fig. S2G-J**). Taken together, these data show that the early pulse regimen of CUT&TIME is a robust and effective method to capture the historical chromatin landscape of RGCs.

### Extended m6A labelling captures the transition between progenitor and RGC

As RGCs differentiate asynchronously and are continuously produced during early retinal development, we reasoned that during a pulse of m6A-MTase expression, m6A labelling becomes enriched at accessible cell-specific DNA elements as a function of the relative amount of time a cell spends in each state. Cells that were RGCs for the entirety of the labeling regimen should show strong RGC chromatin signatures and cells that were progenitors for the entirety of the labelling should show strong progenitor signatures. However, cells that transitioned from progenitor cells to RGCs during m6a-MTase expression would be expected to show a mixture of chromatin states of both progenitor and RGC identity reflective of the relative amount of time the cell spent in each state. Short labeling periods would be expected to show fewer transitioning cells whereas longer labelling periods would be expected to enrich for transitioning cells.

To test this hypothesis, we pulsed the m6a-MTase for 6 days from day 39 to day 45 of retinal development (**Fig. 4A**) and generated a UMAP from these m6a profiles that contained three clusters (**Fig. 4B**). Using progenitor and RGC gene score modules from **Fig. 2**, we were able to identify an RGC cluster, a progenitor cluster, and a cluster with a mixture of progenitor and RGC chromatin signatures which we termed the transition cluster (**Fig. 4C-D**). We revisited our 2-day pulse experiment, and sought to identify similar transition cells that shared RGC and progenitor signatures. We identified a cluster of cells that had intermediate values of both RGC and progenitor module scores, and assigned them to a transition cluster (**Fig. 4E, S3A-B**). While only 14% (727/5246) of cells in the 2-day pulse showed chromatin signatures consistent with cells transitioning from progenitor to RGC, 46% of cells (1618/3491) in the 6-day pulse occupied this transition state (**Fig. 4F**, p < 0.0001 chi-squared test). These trends can also be observed when viewing the gene module score as a violin plot, summarizing the average gene module score for all cells in each cluster for the extended pulse (**Fig. 4G-H**). As transitioning cells capture snapshots of the past and the present chromatin structure, longer 6-day pulse regimen represents a potential alternative strategy to identify terminal differentiation states in cases where FACS sorting is not a viable option.

**Figure 4:**
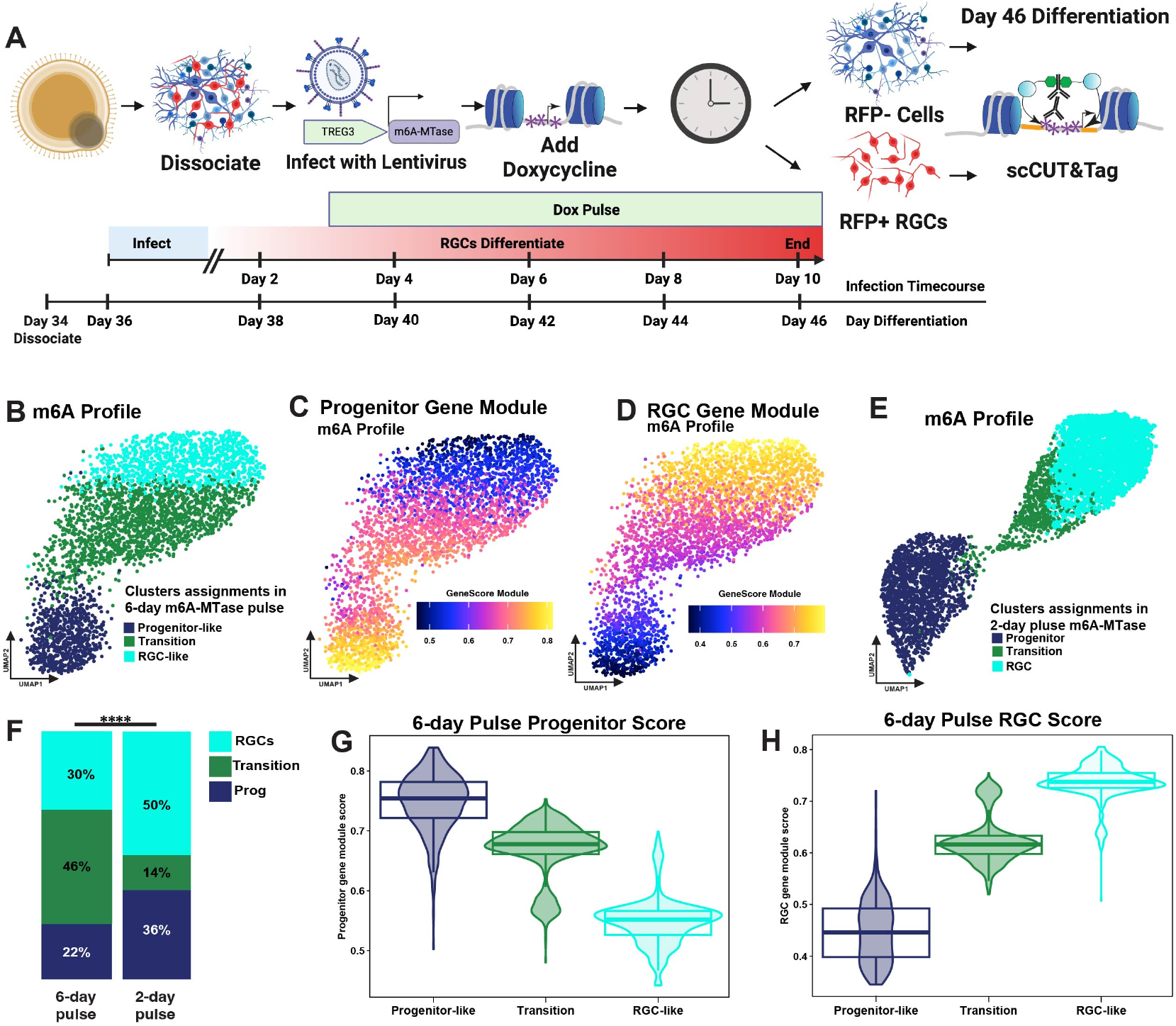
Extended m6A labelling captures transition between progenitor and RGC. **(A)** Schematic of experimental design. **(B-D)** UMAP of m6A profiles generated by CUT&TIME for retinal cells cultured with experimental protocol displayed in **Fig. 4A. (B)** Three clusters defined as progenitor like (blue), transition (green), and RGC-like (cyan). **(C)** Progenitor gene module displaying the m6A profile at progenitor genes displayed in **Fig. 2H. (D)** RGC gene module displaying the m6A profile at RGC genes displayed in **Fig. 2I. (E)** UMAP of m6A profiles generated by CUT&TIME for retinal cells cultured with experimental protocol displayed in **Fig. 2A**.Three different clusters are displayed, progenitor (blue), transition (green), and RGC (cyan). **(F)** Graph showing the percentage of cells in each cluster for the 2-day pulse UMAP displayed in **Fig. 4B** (left, 6-day pulse) and the 6-day pulse UMAP displayed in **Fig. 4E** (right, 2-day pulse), p < 0.0001 chi-squared test. **(G)** Violin plot summarizing the average progenitor gene module score for cells in each cluster shown in **Fig. 4B. (H)** Violin plot summarizing the average RGC gene module score for cels in each sample shown in **Fig. 4B**.

### CUT&TIME identifies promoters associated with the transition from progenitor to RGC

We next sought to use CUT&TIME data to identify genes involved in the transition from progenitors to RGCs. As CUT&TIME allows us to capture the chromatin landscape of the subset of progenitors destined to differentiate into RGCs, we reasoned that differential analysis may allow us to identify changes in chromatin structure that are enriched in progenitors that later differentiate into RGCs as compared to those of self-renewing progenitors.

To perform differential analysis, we grouped our CUT&TIME datasets into three distinct groups: late labeled progenitors, early labeled RGCs (referred to as the “transition” group), and late labeled RGCs. We then ranked each promoter in the human genome according to m6A signal (**Table S1**) and compared the log fold change in rank between groups to obtain the top differentially ranked promoters from each group in our CUT&TIME dataset (**Fig. S4A-C, Table S2**). The progenitor group promoter list included expected markers such as NOTCH1, HES5 and HES4 while the RGC group promoter list included expected markers such as POU4F2, ISL1 and SNCG. The transition promoter list included both known progenitor genes such as SIX3, GLI1, NFIX, NEUROD2, as well as known RGC genes such as KLF16 and TYRO3.

As there is not a perfect correlation between promoter accessibility and gene expression, we sought to narrow the promoter list to genes that were biologically relevant in the retina. Using single cell RNA-seq data from the fetal retina^18^, we narrowed the promoter list to genes that showed differential RNA expression between progenitors, neurogenic precursors (referred to as the transition group), and RGCs (**Table S3**). To assess where in the developmental trajectory these genes show peak accessibility, we applied gene score modules built from each group’s set of enriched and expressed genes. As expected, we see that the genes identified as highly enriched in progenitors are highly represented in the progenitor group (**Fig. 5A, S4D**), and those enriched in RGCs label the RGC group (**Fig. 5B, S4F**). Interestingly, the genes enriched in the transition group show elevated m6A signal primarily in the progenitors, but with elevated accessibility also detectable in the early stages of RGC differentiation (**Fig. 5C, S4E**). This indicates the set of transition genes are initially accessible in progenitors and maintain their accessibility as the cell transitions to becoming an RGC. We see similar m6A enrichment patterns across all CUT&TIME runs when observing gene score modules built from progenitor, transition, and RGCs gene lists (**Fig. S4G-X**), demonstrating that CUT&TIME identifies unique changes in promoters associated with progenitor and RGC identity as well as the transition from progenitor to RGC.

**Figure 5:**
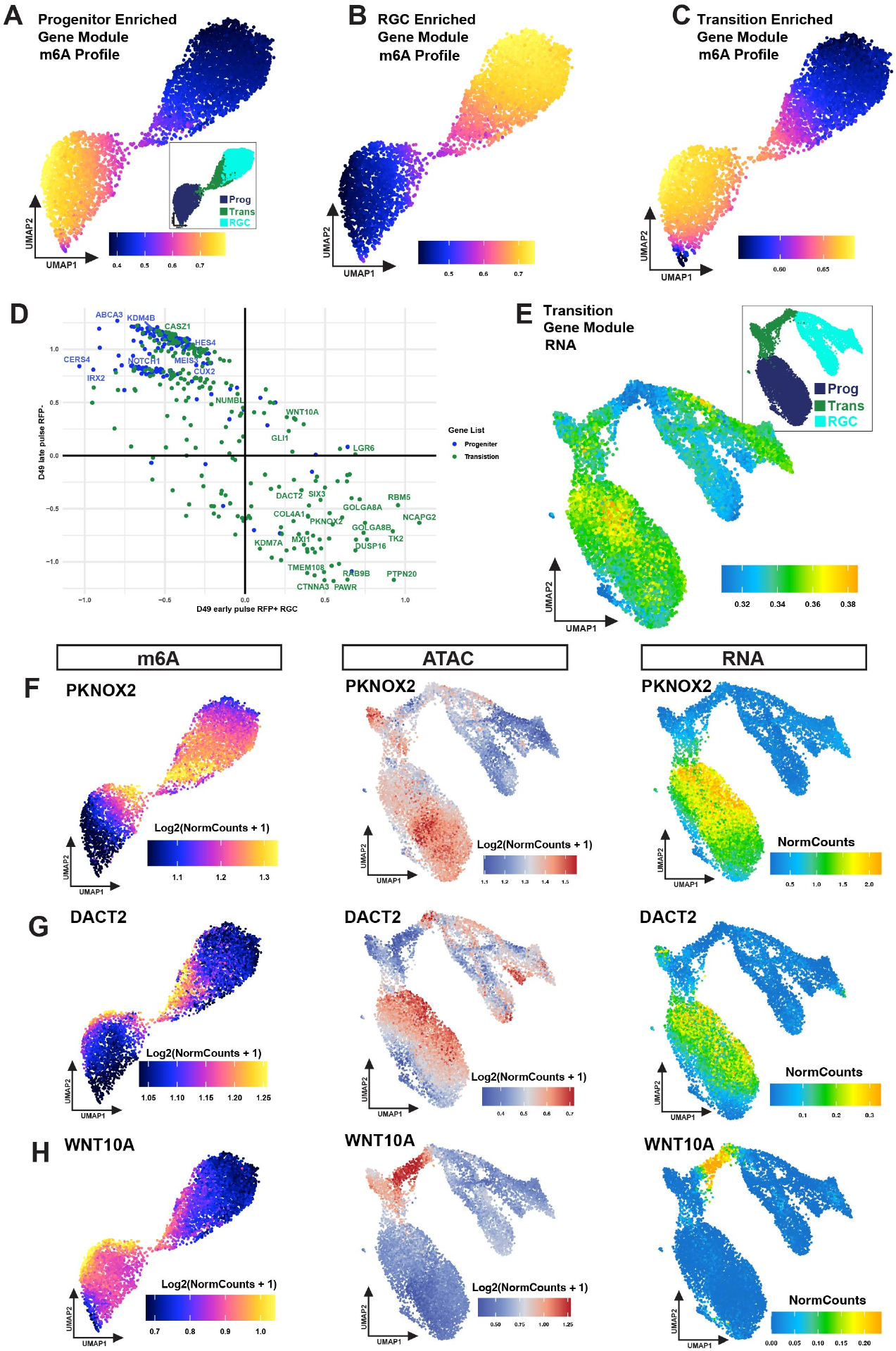
CUT&TIME identifies genes involved in RGC development. **(A-C)** UMAP of m6A profiles generated by CUT&TIME from experimental protocol displayed in **Fig. 2A.** Inset displays three different clusters; progenitor-like (Prog, blue), transition (Trans, green), and RGC-like (RGC, cyan). **(A)** Gene module of progenitor enriched genes listed in **Table S3**. Inset displays three different clusters; progenitor (Prog, blue), transition (Trans, green), and RGCs (RGC, cyan). **(B)** Gene module of RGC enriched genes listed in **Table S3. (C)** Gene module of transition enriched genes listed in **Table S3. (D)** Scatter plot of gene scores for progenitor genes in the D49 late pulse RFP-sample (blue) versus gene scores for transition genes in the D49 early pulse RFP+ RGC (green) from **Table S3. (E)** Gene score module of expressed RNA for all genes from **Fig. 5D** that have a transition gene score that is greater than that of the gene score in the progenitor group applied to a UMAP of multiome data generated previously in the Reh lab from day 59 and day 76 fetal retina^18^. Inset displays three different clusters; progenitor (Prog, blue), transition (Trans, green), and RGCs (RGC, cyan). **(F-H)** UMAP of m6A profiles generated by CUT&TIME from experimental protocol displayed in **Fig. 2A** and **Fig. 5A** for specified genes (left). UMAP of ATAC profiles generated by multiome data displayed in **Fig. 5E** for specified genes (center). UMAP of RNA expression generated by multiome data displayed in **Fig. 5E** for specified genes (right). **(F)** PKNOX2 **(G)** DACT2 **(H)** WNT10A.

We were intrigued that our differential analysis identified distinct genes in the transition group when compared to progenitors, suggesting that distinct chromatin structures and gene expression signatures may distinguish self-renewing progenitors from those destined for differentiation. To better understand the accessibility and expression patterns of candidate genes identified in our transition group, we filtered our transition gene list to contain only those genes with a higher gene score for the transition group compared to the progenitor group (**Fig. 3D, Table S4**). We then used this modified transition gene list to create another gene module, which was applied to the fetal retina UMAP of RNA expression (**Fig. 3E**). We observe an enrichment for RNA expression in a subset of progenitors, indicating that many of these genes maybe variably expressed in a subpopulation of progenitors. Further investigation of individual candidate genes revealed genes showing high expression in a subpopulation of progenitor cells. One such candidate was PKNOX2, a transcription factor belonging to the three amino acid loop extension (TALE) class of homeodomain proteins. PKNOX2 has been reported to play an important role in mammalian cochlea development^19^, and has also been shown to be a tumor suppressor gene in gastric cancer^20^. ATAC-seq and RNA-seq data indicate that PKNOX2 is expressed in a subpopulation of retinal progenitors (blue cluster, “prog”), similar to what is observed for CUT&TIME data (**Fig. 5E, F**). PKNOX2’s expression in a subset of progenitors together with its identification as a transition gene in our CUT&TIME data suggest that PKNOX2 may mark a subset of progenitors preparing to differentiate. However, a functional role remains unclear, as loss of function of PKNOX2 in the retina has not yet been tested.

We also identified DACT2, a binding antagonist of β-catenin that has been shown in several cancer contexts to block proliferation by inhibiting Wnt/β-catenin signaling^21^. Loss of DACT2 in zebrafish has been shown to cause reduced eye size^22^, suggesting that DACT2 may also regulate cell proliferation in the developing retina. With CUT&TIME, we find this gene is labeled in progenitors that transition to become RGCs, indicating that it may be important for influencing the transition from progenitor identity to RGC. When querying DACT2 expression in the developing retina, we see it is weakly expressed in a sub-population of progenitor cells, similar to CUT&TIME (**Fig. 5G**). Together, these data suggest that DACT2 may act to block progenitor proliferation as the cell is transitioning towards becoming a retinal ganglion cell.

Finally, we identified WNT10A as a candidate gene from the CUT&TIME analysis. WNT signaling has been previously shown to play a critical role in retinal development^23,24^, but a role for WNT10A has not been explicitly tested. Multiome analysis indicates that WNT10A gains accessibility and is transiently expressed exclusively during the transition from progenitor to RGC during retinal development (cyan cluster, “RGC”) (**Fig. 5E, H**). CUT&TIME, however, indicates that WNT10A gains m6A accessibility in a subset of progenitors prior to the RGC transition. These data suggest that WNT10A expression in the transition zone may be primed in a limited population of progenitors dedicated to transitioning to RGCs. As these early changes in accessibility are undetectable in the progenitor cluster with ATAC-seq, these data demonstrate that CUT&TIME is capable of detecting changes in chromatin accessibility that precede detectable changes by ATAC-seq.

Together, these data reveal unique expression patterns of genes in the transition group and identify candidates for future investigations to understand their roles in RGC specification.

## Discussion

A fundamental question in developmental biology concerns identifying the earliest changes in progenitor chromatin structure associated with cell fate commitment. Here we present CUT&TIME, a new technique that generates a record of the open chromatin in living cells, that can be read out at later time points in development. By generating a record of past chromatin accessibility, CUT&TIME provides a means to directly visualize the chromatin structure of progenitor cells that gave rise to specific cell types during differentiation. By comparing the chromatin landscape of progenitor cells destined to differentiate into RGCs to all progenitor cells, we demonstrate that we are able to identify distinct chromatin structures in progenitor cells that are associated with cell fate commitment.

CUT&TIME builds upon previous methods that have employed m6A labelling to mark open chromatin regions^11^, or to mark DNA associated with certain chromatin features^9,25^. m6A labeling is compatible with living cells, and other methods have leveraged this feature to capture dynamic changes in lamin-associated DNA^8^ and RNA polymerase II^9^. However, as these methods are targeted to specific chromatin features or require bulk sequencing approaches, they are limited in their ability to capture global changes in chromatin accessibility at the single cell level. By combining untethered m6A-MTase labeling with single cell sciCUT&Tag profiling, CUT&TIME gives an unbiased view of the past open chromatin landscape on a single cell level, which for the first time enables interrogation of the diversity of chromatin states that give rise to a particular cell fate during differentiation.

Much of our current understanding of changes in chromatin structure during development has been derived from pseudotime trajectories built from single cell-compatible modalities such as ATAC-seq. Pseudotime trajectories are built by taking a single snapshot in time of the gene expression or chromatin accessibility landscape and then calculating the most probable path of development based on transcriptomic similarities between the existing cells. While powerful, these techniques are built upon inferences, and cannot directly reconstruct the past chromatin state of differentiated cells. Furthermore, differences in chromatin structure can only be inferred to pertain to cell-fate specification if they occur in transition zones, which refer to regions in UMAP space that separate two cell states, such as progenitors and RGCs. Differences in chromatin structure are also detectable amongst cells uniformly classified as progenitors (**Fig. 5E**). However, using pseudotime analysis, it is challenging to separate early changes in chromatin structure associated with differentiation from fluctuations in gene activity unrelated to cell fate commitment. By directly tracking the past chromatin structure of differentiated cells over multiple experiments, CUT&TIME can identify changes in progenitor chromatin structure which are consistently associated with progenitors that undergo cell fate specification. In this way, CUT&TIME can help identify the earliest changes in progenitor chromatin structure that shift a progenitor from a program of self-renewal to differentiation.

By comparing all retinal progenitors to those progenitors which differentiated into RGCs, we have identified a group of genes that show dynamic changes in chromatin structure during progenitor differentiation that may be important for the development of progenitors to RGCs. We hope to utilize this technique in future studies to profile other terminal cell types generated in the retina to identify which genes are necessary specifically for RGC development, and which genes are necessary more generally for retinal cell fate specification. This will make it possible to identify the earliest developmental decisions that progenitors undergo as they specify the diversity of cell types within the retina.

In the future, it would be exciting to combine CUT&TIME with current lineage tracing methods involving DNA editing that can identify lineage relationships in developmental systems via DNA sequencing^26-31^. These lineage tracing methods provided valuable insights into developmental trajectories, but do not assess global changes in chromatin structure that specify distinct cell fates. By pairing these technologies with CUT&TIME, we could understand how chromatin structure shifts in a single lineage over developmental time to generate the diversity of cell types needed to pattern the mature retina.

While CUT&TIME is effective in capturing and preserving chromatin structure over time scales as long as 12 days, the technology remains limited by the fact that mammalian cells do not have the machinery to propagate the m6A mark onto newly synthesized DNA during cell division. This means that over multiple cell divisions the mark will be diluted, making it much harder to read out the history of chromatin structure in systems that are highly proliferative. To overcome this limitation it may be possible to express the bacterial enzymes that carry the mark forward during replication^32^, and thereby mark the lineage of a cell through multiple divisions. This would allow future studies to trace the chromatin history of retinal cells specified at the end of development, such as rods and Müller glia.

CUT&TIME has identified a group of genes that undergo dynamic changes in structure and may be important for the transition of progenitors to RGCs. This technology is widely applicable to other model systems to better understand how dynamic chromatin structures shape cell fate decisions in complex tissues during development, homeostasis, neuronal plasticity, and regeneration following tissue damage or injury.

## Acknowledgements

We would like to thank all the members of the Reh, Henikoff, Stergachis, and Bermingham-McDonogh labs for their valuable comments on the manuscript. We thank Don Zack for his generosity in sharing the WA07 Brn3b-tdTomato hESC line, and David Gamm for his generosity in sharing the WA09 NRL+/eGFP hESC line. This research was supported by Foundation Fighting Blind-ness (TA-RM-0620-0788-463 UWA to T.A.R.), a gift from Open Philanthropy (to T.A.R), the Damon Runyon Cancer Research Foundation (DRG-# 32-20 to K.C.E.), the National Institutes of Health (NIH) LRP (to K.C.E.), and a Hanna H. Gray Fellows Program Award from the Howard Hughes Medical Institute (Grant #GT15994 to K.C.E.), the Howard Hughes Medical Institute (to S.H.), a K99 pathway to independence award, NIGMS (Grant #GM152821-02 to M.W.), University of Washington Genome Training Grant (to M.W.), NIH grant 1DP5OD029630 (to A.B.S.), a Career Award for Medical Scientists from the Burroughs Wellcome Fund (to A.B.S.), and A.B.S. is a Pew Biomedi-cal Scholar. Research reported in this publication was generated using the DLMP Flow Cytometry Core, and stem cell lines were generated in part through the Tom and Sue Ellison Stem Cell Core at the University of Washington. Illustrations were created with BioRender.com.

## Author contributions

Conceptualization, K.C.E, M.W., T.A.R., and S.H.; Methodology, K.C.E, and M.W.; Software, K.C.E, M.W., and S.J.N.; Formal Analysis K.C.E, and M.W. J.H.; Investigation, K.C.E, M.W., D.J.J., S.J.N., J.E.R., S.J.E., J.W., S.M.S., G.W.D.; Resources, S.H., T.A.R., A.B.S.; Writing – Original Draft, K.C.E.; Writing –Review & Editing, K.C.E, M.W., T.A.R., S.H.; Funding Acquisition, T.A.R., K.C.E., S.H., M.W., A.B.S., Supervision, K.C.E, M.W., T.A.R., S.H.

## Supplemental information

**Document S1**. Figures S1–S4 and Table S5.

**Table S1:** Table_S1.xlsx, excel file containing the ranked promotors for each data set included in the analysis, related to Figure 5.

**Table S2:** Table_S2.xls, excel file containing the top differentially ranked promoters for progenitors, the transition group, and RGCs in our CUT&TIME dataset pertaining to Fig. 5 and displayed in Fig. S4A-C.

**Table S3:** Table_S3.xls, excel file containing the top differentially accessible promotors from Table S2 that are expressed in 59 human fetal multiome data used to create the progenitor enriched, RGC enriched, and transition enriched gene modules, related to Figure 5.

**Table S4:** Table_S3.csv, csv file containing gene scores for progenitor genes in the D49 late pulse RFP-sample versus gene scores for transition genes in the D49 early pulse RFP+ RGC from Table S3, related to Figure 5.

**Table S5**

## Materials and Methods

### Stem cell and organoid culture

hESCs or hiPSCs were maintained by dissociation and propagation in StemFlex media (ThermoFisher, A3349401) on Matrigel (Corning, CLS354277) coated plates at 37 C in 5% CO2. hESC or hiPSC colonies were passaged by dissociation with ReLeSR (StemCell Technologies, 100–0484). Retinal organoids were grown as described in Tresenrider et al., 2023^33^, with minor alterations mentioned herein. The cell culture media used were neural induction media (NIM): 484.5mL DMEM/F12 (Life Technologies, Catalog #11330-057), 5 mL N2 supplement (Life Technologies, Catalog #17502048), 5 mL MEM NEAA (Life Technologies, Catalog# 11140050), 5 mL penicillin-streptomycin (Life Technologies, Catalog# 15240062). Retinal differentiation media (RDM): 240 mL DMEM/F12 (Life Technologies, Catalog #11330-057), 240mL DMEM (Life Technologies, Catalog #12430062), 10 mL B27 supplement (Life Technologies, Catalog #17504001), 5 mL MEM NEAA (Life Technologies, Catalog #11140050), 5 mL penicillin-streptomycin (Life Technologies, Catalog #15240062) + 1%, or 10% FBS (Corning, Catalog #35-011-CV) based on protocol needs.

Retinal organoid culture protocol was adapted from Tresenrider et al., 2023^33^. In brief, confluent stem cell colonies were lifted from the plate using dispase (2 mg/mL, Life Technologies, Catalog #17105041) for 5–10 min, then forcefully removed by pipetting 2 mL/well of DMEM (Life Technologies, Catalog #12430062) directly onto cells using a 1 mL pipette. Colonies were removed from the 6 well plates and allowed to settle to the bottom of a 15 mL collection tube by gravity. The supernatant was removed, and colonies were transferred to a T25 tissue culture flask in a 1:1 StemFlex:NIM mix for a total of 10 mL per flask. Full media change was performed the following day (Day 1) with 10 mL of NIM. Day 2-5 a full media change was performed and replaced with 10 mL of NIM. Day 6 a full media change was performed and re-placed with 10 mL of NIM, and BMP4 (R&D Systems, 314-BP-050) was added to the media to a final concentration of 1.5nM. Day 8 EBs were then evenly distributed between the 6 wells of a 6 well plate in 2mLs of fresh NIM. 200 uL of FBS was added to each well to allow for EBs to stick to the bottom of the plate. Days 10–20, every other day, a full media change was performed with fresh NIM. Days 14 and 16 a full media change was performed with NIM, and the signaling factor modulator CHIR99021 (Cedarlane Laboratories, 04-0004-02) was added to the media to a final concentration of 3mM. EBs were then manually lifted from the plates on day 20 or 21 by forcefully pipetting 2 mL/well of DMEM directly onto EBs using a 1 mL pipette. EBs were removed from the 6 well plates and allowed to settle to the bottom of a 15 mL collection tube by gravity. From this point on we consider the cells retinal organoids. The supernatant was removed, then organoids were resuspended in RDM 10% FBS. From this point on, cells were maintained with a full media change of RDM +10% FBS every two to three days.

### Stem cell lines used

The cell lines we use here are the following: WA09 NRL+/eGFP hESC line generated by the Gamm lab^34^, and the WA07 Treg Brn3b-tdTomato reporter hESC line developed as described below.

WA07 Treg Brn3b-tdTomato reporter hESC line generation: This line was originally developed in the Zack lab to insert the Brn3b-tdTomato reporter into WA07 ESCs^35^. WA07 Brn3b-tdTomato reporter hESCs were cultured on Matrigel-coated plates in mTeSR1 media and passaged using Accutase. The UW Tom & Sue Ellison Stem Cell Core then generated an AAVS1-TRE3-Dicer construct by cloning Dicer cDNA containing a silent mutation that disrupts the PAM sequence of Dicer cr6 gRNA^36^ into the pAAVS1-TRE3-GFP construct flanked with AAVS1 homology arms (Addgene #52343) using Gibson cloning.

One million WA07 Brn3b-tdTomato reporter hESCs were electroporated with the resulting AAVS1-TRE3-Dicer construct (4ug) and plasmids expressing TALEN targeting the AAVS1 locus (TALEN-R and TALEN-F, Addgene #52341and 52342; 0.5ug of each) using Amaxa Human Stem Cell Nucleofector (kit 2 Lonza). Two days following the nucleofection, the cells were selected with puromycin 0.5µg/ml for 2 days. Individual colonies were hand-picked and plated into 96 well plates. Genomic DNA was extracted using Quick Extract DNA extraction solution (Epicentre#QE09050) and clones were screened by PCR using Phusion High Fidelity PCR master mix (Thermo Fisher Scientific).

Genotyping primers utilized to detect positive clones:

Reaction #1, iDicer insertion: F (iDicer6m-f) AGTTGGTTGCACGGG-TATTTCC, R (LNCX-r)-GCTCGTTTAGTGAACCGTCAGATC (3.7kb);

Reaction #2, Insertion into the AAVS1 locus: F(dna803)-TCGACTTCCCCTCTTCCGATG, R(dna804)-GAGCCTAGGGCCGGGAT-TCTC (1.2kb);

Reaction #3, mono-vs bi-allelic insertion: F(dna803)-TCGACTTCCCCTCTTCCGATG, R(dna183)-CTCAGGTTCTGGGA-GAGGGTAG (1.4kb).

These cells have the Treg inducible system that activates the inducible m6A-MTase on the lentiviral vector. There is also an exogenous DICER allele that is expressed upon Dox addition, however this has not been seen to have any effect on retinal development or cell type differentiation according to histological examination.

### Tissue dissociation for plating and infection

Organoids were selected for visually, or for containing Brn3b-tdTomato+ cells, then dissociated using Accutase and DNAse for 15-20 minutes on a nutator at 37C, with gentile pipetting to break up cells every 7 minutes. Cell pellets were resuspended in RDM + 1% FCS and plated onto a Matrigel coated well. Cells were cultured for 12 – 15 days. Lentiviral infection was carried out on day 1 or 2 of plating. For infection, 2 ug/mL Polybrene with variable titers of virus obtained through Vector Builder were added to RDM + 1% FCS for 48 hours, then removed. If doxycycline was needed for induction of m6A-Mtase expression, 3 ug/mL was added to the cultures every two days for a total of 2 or 4 days of doxycycline exposure.

### Tissue dissociation for CUT&TIME and Fiber-seq

Plated cells were washed with DBPS and then dissociated from the plate with Accutase and DNAse for 7-14 minutes on a nutator at 37C, with gentile pipetting to break up cells every 7 minutes. Cell pellets were resuspended in RDM + 1% FCS and placed on ice. If required for the experiment, cells were then sorted on the BD FACSAria III sorter based on RFP+ or RFP-signal at the DLMP Flow Cytometry Core.

### CUT&TIME methodology

Organoid cells were pelleted and resuspend in 1 ml Wash Buffer1 (20 mM HEPES at pH 7.5, 150 mM NaCl, 0.5 mM spermidine, and 1× protease inhibitors) + 0.05% digitonin. Cells were then placed on ice while beads were prepped. 100 ul of bangs 5-micron beads were activated and added to cells, then incubated for 10 min on ice. The supernatant was then removed, and bead-bound cells were resuspended in 100 ul of Wash Buffer1 + 0.05% digitonin. Cells were then transferred to PCR tubes in preparation for heating. Supernatant was removed again and cells were resuspended in 50 heating buffer (1ml Wash Buffer1 + 0.05% digitonin + 1 ul EDTA) and transferred to PCR machine. Samples were then heated at 98°C for 20 minutes. Samples were then removed from PCR machine and allowed to cool at room temperature for 15 seconds. 1 ul of anti-m6A antibody was then added and samples were mixed by gently pipetting. Samples were allowed to cool for an additional minute, after which 1 ul of anti-m6A antibody was added and samples were mixed with gentle pipetting. Samples were allowed to cool for an additional 2 minutes, after which 1 ul of anti-m6A antibody was added and samples were mixed with gentle pipetting. Samples were allowed to cool for an additional 5 minutes, after which 1 ul of anti-m6A antibody was added and samples were mixed with gentle pipetting. Samples were then placed at room temperature for 10 minutes, after which, 200 uls of ice cold Wash Buffer1 + 0.05% digitonin were then added. Samples were then placed on magnet and supernatant removed. Samples were then washed twice with 200 uls Wash Buffer1 + 0.05% digitonin. Supernatant was again cleared and cells were resuspended in 250 uls Wash Buffer1 + 0.05% digitonin + 1.6 ul 16% formaldehyde. Samples were mixed by gentle pipetting and placed on ice for two minutes. Samples were then placed on magnet and supernatant removed. Samples were then resuspend in 50 uls antibody binding buffer (1 ml Wash Buffer1 + 0.05% Triton + 4 uls EDTA) with 1 ul anti-6mA antibody. Samples were allowed to incubate overnight, after which samples were processed according to standard sciCUT&Tag protocol as previously described in Janssens et al., 2024^37^.

### CUT&Tag methodology

CUT&Tag reactions were performed according to the CUT&Tag PCI protocols Wooten et al. 2023^38^, and Kaya-Okur et al. 2020^39^, with some modifications. Briefly, ∼100,000 cells were spun down and resuspended in Wash Buffer1 (20 mM HEPES pH 7.5, 150 mM NaCl, 0.5 mM spermidine, 0.05% Triton-X100, Roche Complete Protease Inhibitor EDTA-Free tablet) by gentle pipetting. 5 ul of Bio-Mag Plus Concanavalin A (ConA)-coated magnetic beads (Bangs Laboratories Catalog #BP531) per sample were activated and added to the cells, then incubated for 10 min on ice. ConA-bound cells were suspended in antibody binding buffer (Wash Buffer1 containing 2 mM EDTA) and split into individual 0.5 ml tubes for overnight incubation with the primary antibody at a 1:100 dilution at 4°C. Samples were placed on magnet and supernatant removed followed by 3x wash with Wash Buffer1. Samples were then resuspended in Wash Buffer1 containing the secondary antibody (guinea pig anti-rabbit IgG) at a 1:50 dilution and incubated at 4°C for 1 h. Following another wash, the samples were resuspended in 300-wash buffer (Wash Buffer1 with an additional 150 mM NaCl) containing Protein AG-Tn5 (pAG-Tn5 at a 1:20 dilution, EpiCypher, Catalog #15-1117) and incubated at 4°C for 1 h. The samples were then washed in 300-wash buffer and resuspended in tagmentation buffer (300-wash buffer with 10 mM MgCl2), followed by incubation at 37°C for 1 h to complete the Tn5 tagmentation reaction, after which the DNA was extracted via phenol-chloroform isoamyl alcohol (PCI) for library preparation. Twenty-one microliters of DNA was mixed with a universal i5 and a uniquely barcoded i7 primer and amplified with NEB Q5 high-fidelity 2× master mix (catalog no. M0492S). The libraries were purified with 1.1× volume of Sera-Mag carboxylate-modified magnetic beads and subjected to LabChip DNA analysis and Illumina sequencing.

### Fiber-seq methodology

Fiber-seq protocol was performed as described in Stergachis et al., 2020^11^ with minor alterations. In brief, nuclei from sorted RFP+ or RFP-cells were isolated as previously described in Thurman et al., 2012^40^ MTase reaction was performed using 0.5 µl of Hia5/million cells. The MTase reactions were incubated for 10 minutes at 25 C, then stopped with 3 µl of 20% SDS (1% final) and transferred to new 1.5 mL microfuge tubes. The sample volumes were increased by adding 130 µl Buffer A and an additional 7 µl of 20% SDS. All samples were mixed with 2 µl RNase A (Invitrogen AM2271) and 2 µl Proteinase K (NEB P8107S) and incubated for 1 hour at 50°C. The DNA was purified using the Promega Wizard kit (Promgega A2920). DNA was then re-purified using AMPure beads, quantified using a Fragment Analyzer (Agilent) and then sheared using a Megaruptor (Diagenode Diagnostics) to 10 15kb in length. The sheared sample was used to generate a Pacific Biosciences Sequencing library using the SMRTbell Prep kit by the University of Washington Long Reads Sequencing Center (Seattle, WA, USA), and libraries were prepared and sequenced samples were pooled and sequenced on two SMRT cells.

### Fiber-seq analysis

Fibertools was used to identify base-pair resolution m6A marks and broader methylation-sensitive patches (MSPs) along individual DNA molecules^41^. Annotated reads were subsequently aligned to the human reference genome (hg19), and regions of aggregated accessibility were detected using the FIRE algorithm^42^.

### Promoter Rank Fold Change Analysis

To generate plots shown in Fig. S4A-C comparing promoter rank change across groups, the computeMatrix function of Galaxy was used to generate coverage plots of normalized read counts of a 10 kb window centered on each promoter in the human genome. The resulting matrices generated from this function were then plotted in decending order using the plotHeatmap function of Galaxy with the “save the regions function” selected. This generated a ranked list of all promoters in the human genome ranked according to their relative CUT&TIME coverage with a rank of 1 denoting the highest amount of CUT&TIME coverage. We then created an average rank for all promoters by averaging promoter ranks across all experiments. To generate a rank fold change value for each group, we then divided the rank for each group (late labeled Progenitors, Transition, and late labeled RGCs), by the population average, to generate a fold change in rank. We set a 1.4-fold rank change as the cutoff for promoters to be considered as differing in rank between groups, as this was the approximate inflection point in a knee-plot of transition group rank change *versus* population average. If promoters appeared multiple times in different groups, that promoter was assigned to the group whose rank indicated greater overall coverage. For example, if the late-labelled progenitor rank was 95 and the transition group rank was 125, that promoter would be assigned to the progenitor group. Once these promoter lists were generated for progenitors, transitions, and RGCs, we sought to verify if their gene expression changed significantly using a previously generated human fetal D59 mulitome dataset^18^. The dataset was subsetted to progenitors, neurogenic precursors (to represent transition cells), and RGCs and a Wilcoxon rank sum test was performed using Seurat’s FindAllMarkers command to find genes that where enriched in at least one cell type. A promoter was considered to have significant gene expression differences if its associated gene showed up as differentially expressed with an adjusted p-value of less than.01 in the Wilcoxon rank sum test in at least one cell type.

### Sectioning and immunofluorescence

Organoids were fixed in 4% paraformaldehyde, 1X PBS, and 5% sucrose at 4C for 45 minutes followed by 3 washes in 1X PBS, 5% sucrose. All samples were then transitioned into 10%, 20% and 30% solutions of sucrose. These were then embedded in OCT and cryosectioned at 14-16 mm.

Immunostaining was performed as previously described in Hoshino et al., 2017^43^. Briefly, slides were blocked in 0.5%Triton/2% horse serum for an hour, followed by primary antibody incubation in 0.5%Triton/2% horse serum at 4C overnight. The next day, the slides were washed 3 times in 1X PBS, followed by secondary antibody incubation in 0.5%Triton/2% horse serum for an hour at room temperature then washed three times with PBS before being mounted. Slides were then mounted using Fluromount. Antibodies are listed in **Table S5**.

### Microscopy

Brightfield images were obtained at 4x magnification using the Zeiss Axio Observer D1. Confocal images were obtained using the Zeiss LSM 990. Image processing was done in ImageJ and Adobe Photoshop.

## Data and code availability

The datasets generated within this study will be available on GEO upon publication in per reviewed journal. Single cell analysis was completed in ArchR, Version 1.0.3. Statistics were calculated in RStudio and Prism 10.

## Supplementary Information

**Table S1:** Table_S1.xlsx, excel file containing the ranked promotors for each data set included in the analysis*

**Table S2:** Table_S2.xls, excel file containing the top differentially ranked promoters for progenitors, the transition group, and RGCs in our CUT&TIME dataset pertaining to Fig. 5 and displayed in Fig. S4A-C.*

**Table S3:** Table_S3.xls, excel file containing the top differentially accessible promotors from Table S2 that are expressed in 59 human fetal multiome data used to create the progenitor enriched, RGC enriched, and transition enriched gene modules.*

**Table S4:** Table_S3.csv, csv file containing gene scores for progenitor genes in the D49 late pulse RFP-sample versus gene scores for transition genes in the D49 early pulse RFP+ RGC from Table S3.*

*Available upon publication in peer-reviewed journal, or by corespondance with T.A.R. –

**Table S5.**

| Antibody | Source | Catalog # | Dilution |
| --- | --- | --- | --- |
| OTX2 | R&D Systems | BAF1979 | 1:300 |
| DAPI | Invitrogen | D1306 | 3 mM |
| GFP | ABCAM | AB13970 | 1:1000 |
| CRX | Santa Cruz | sc-377138 | 1:500 |
| Donkey anti-goat 568 | Invitrogen | A11057 | 1:300 |
| Donkey anti-mouse 647 | Invitrogen | A32787 | 1:300 |
| Donkey anti-chicken 488 | Jackson Immuno | 703-545-155 | 1:500 |
| N6-Methyladenosine (m6A) (D9D9W) | Cell Signaling | 56593 | 1:100 |
| anti-RNAPII Ser2-phosphorylation | Cell Signaling | 13499S | 1:10 |
| anti-RNAPII Ser5-phosphorylation | Cell Signaling | 13523S | 1:10 |
| anti-RNAPII Ser5/Ser2-phosphorylation<br>Phospho-Rpb1 CTD (Ser2/Ser5)<br>(D1G3K) | Cell Signaling | 13546S | 1:10 |

**Supplementary Figure 1.**
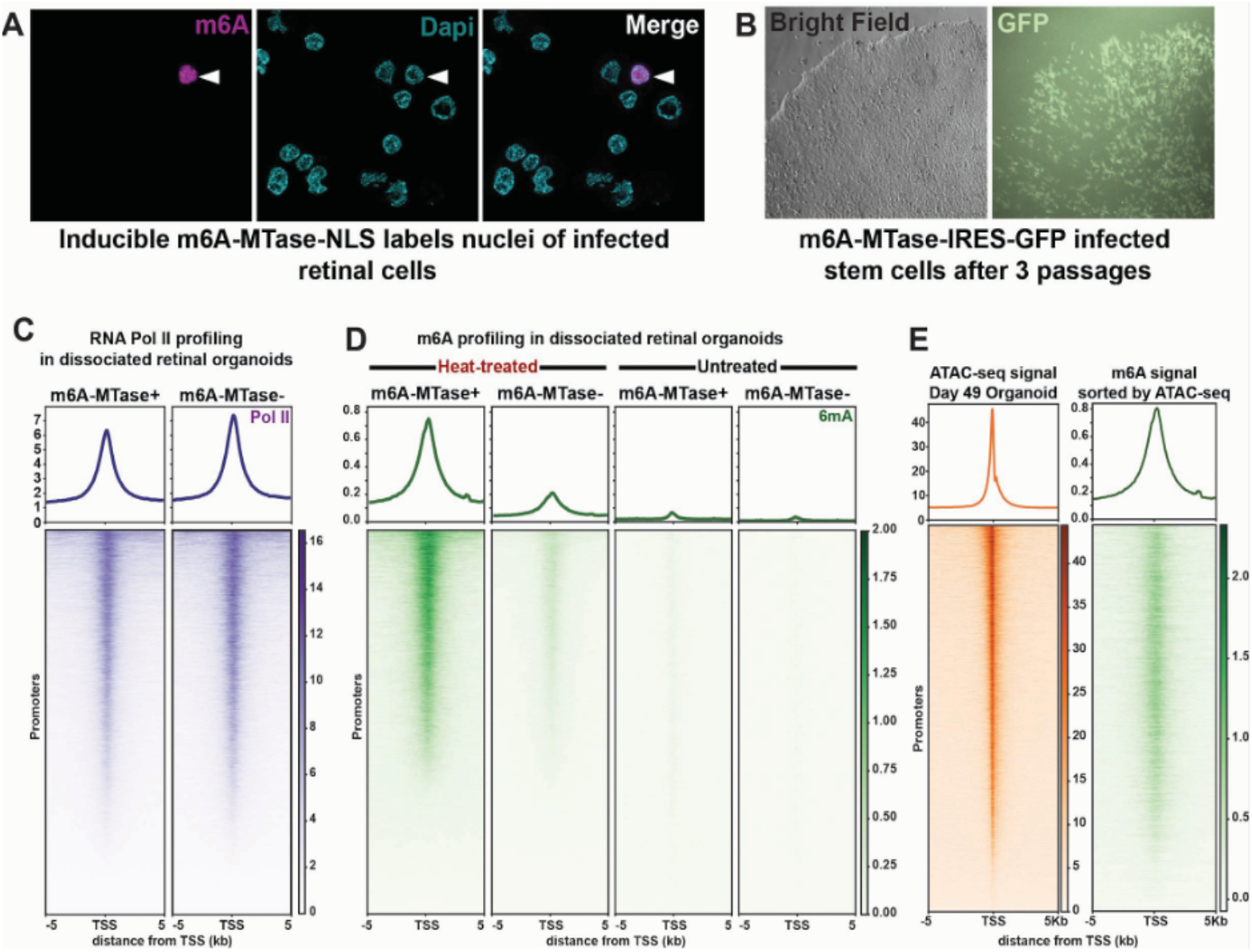
Related to Figure 2. A) Retinal cells infected with m6A-MTase-NLS, stained with m6A antibody (magenta) and DAPI (cyan). B) Stem Cells infected with m6A-MTase-IRES-GFP infected and passaged three times. Bright field on left, GFP on right. C-E) Heatmaps of profiles 5Kb on either side of all promotors is shown, ordered from most coverage to least coverage. Graphs at the top display the average distribution of the signal along the promotors. C) RNA Pol II profiling in dissociated and plated retinal organoids, cultured similarly as displayed in **Fig. 1A**. Left: cells infected with m6A-MTase, Right: uninfected cells. Genes are in the same order in the two plots. D) m6A profiling in dissociated and plated retinal organoids, cultured similarly as displayed in **Fig. 1A**, in four different experimental conditions. Leftmost two columns: samples were heat-treated during the CUT&Tag protocol, and samples were infected with m6A-MTase (left) or uninfected (right). Rightmost two columns: CUT&Tag protocol was performed as normal, and samples were infected with m6A-MTase (left) or uninfected (right). Genes are in the same order in the four plots. E) Left: Heat map of ATAC-seq signal from day 49 retinal organoids previously generated in the Reh lab^16^. Right: m6A profiling from CUT&TIME protocol with promotors presented in the same order as those from the ATAC-seq.

**Supplementary Figure 2.**
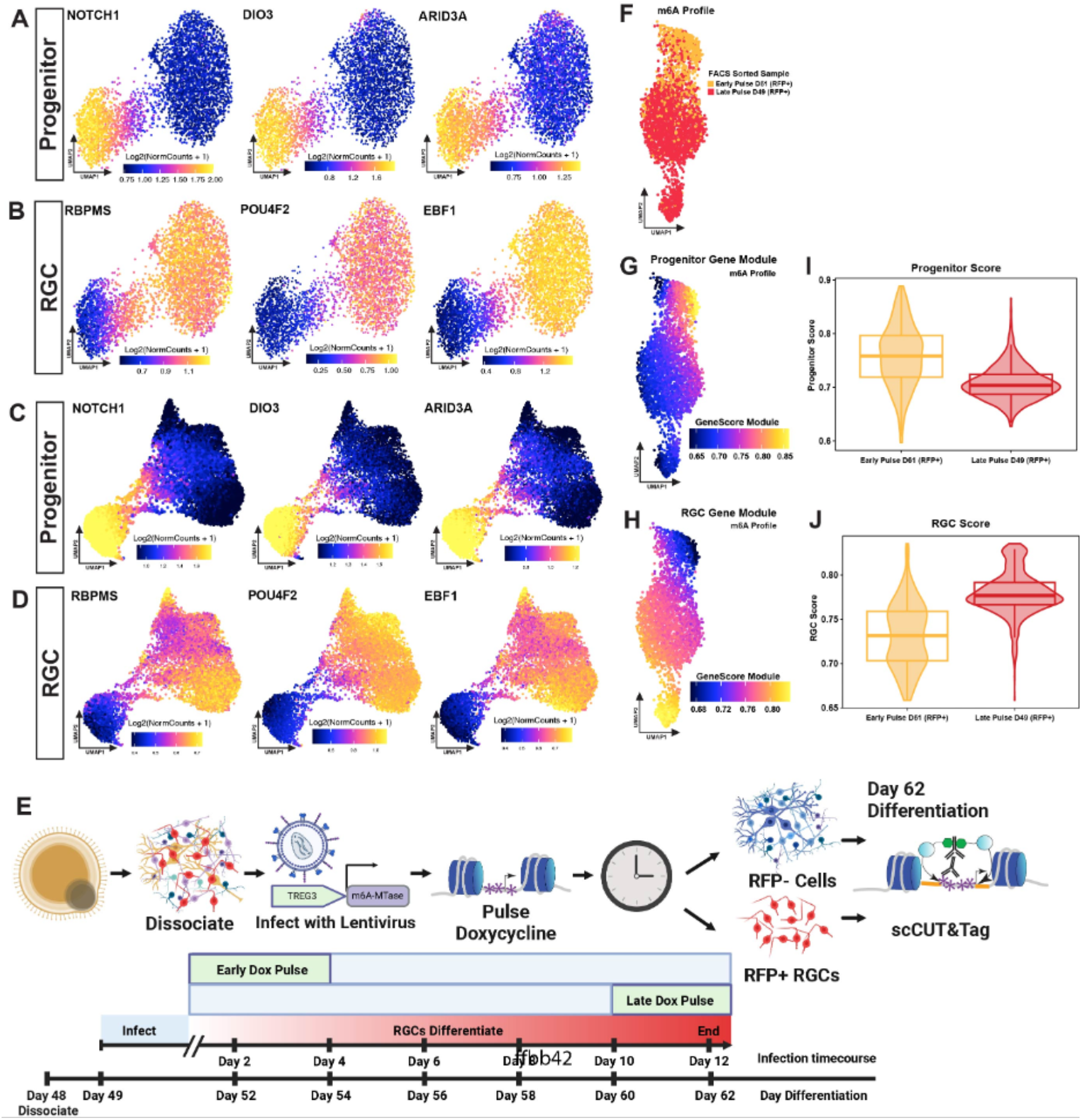
Related to Figure 3. A) UMAPs of RNA Pol II profiles generated by CUT&Tag for retinal cells cultured with experimental protocol displayed in **Fig. 3B**, showing the gene score of profiling density along the progenitor genes NOTCH1, DIO3, and ARID3A. B) UMAPs of RNA Pol II profiles generated by CUT&Tag for retinal cells cultured with experimental protocol displayed in **Fig. 3B**, showing the gene score of profiling density along the RGC genes RBPMS, POU4F2, and EBF1 (bottom). C) UMAPs of m6A profiles generated by CUT&TIME for retinal cells cultured with experimental protocol displayed in **Fig. 3E**, showing the gene score of profiling density along the progenitor genes NOTCH1, DIO3, and ARID3A. D) UMAPs of m6A profiles generated by CUT&TIME for retinal cells cultured with experimental protocol displayed in **Fig. 3E**, showing the gene score of profiling density along the RGC genes RBPMS, POU4F2, and EBF1. E) Schematic of experimental design for early pulse day 61 cluster displayed F-I. F-H) UMAP of m6A profiles generated by CUT&TIME for day 61 early pulse retinal cells cultured with experimental protocol displayed above in E, and day 49 late pulse retinal cells cultured with experimental protocol displayed in **Fig. 2A**. F) UMAP of m6A signal generated by CUT&TIME. Cells were sorted based on RFP signal into two groups: D61 early pulse RFP+ ganglion cells (orange), and D49 late pulse RFP+ cells (red). G) Progenitor gene module displaying the m6A profile at progenitor genes displayed in **Fig. 2H**. H) RGC gene module displaying the m6A profile at RGC genes displayed in **Fig. 2I**. I) Violin plot summarizing the average progenitor gene module score for cells in each sample shown in **Fig. 2D**. J) Violin plot summarizing the average RGC gene module score for cels in each sample shown in **Fig. 2D**.

**Supplementary Figure 3.**
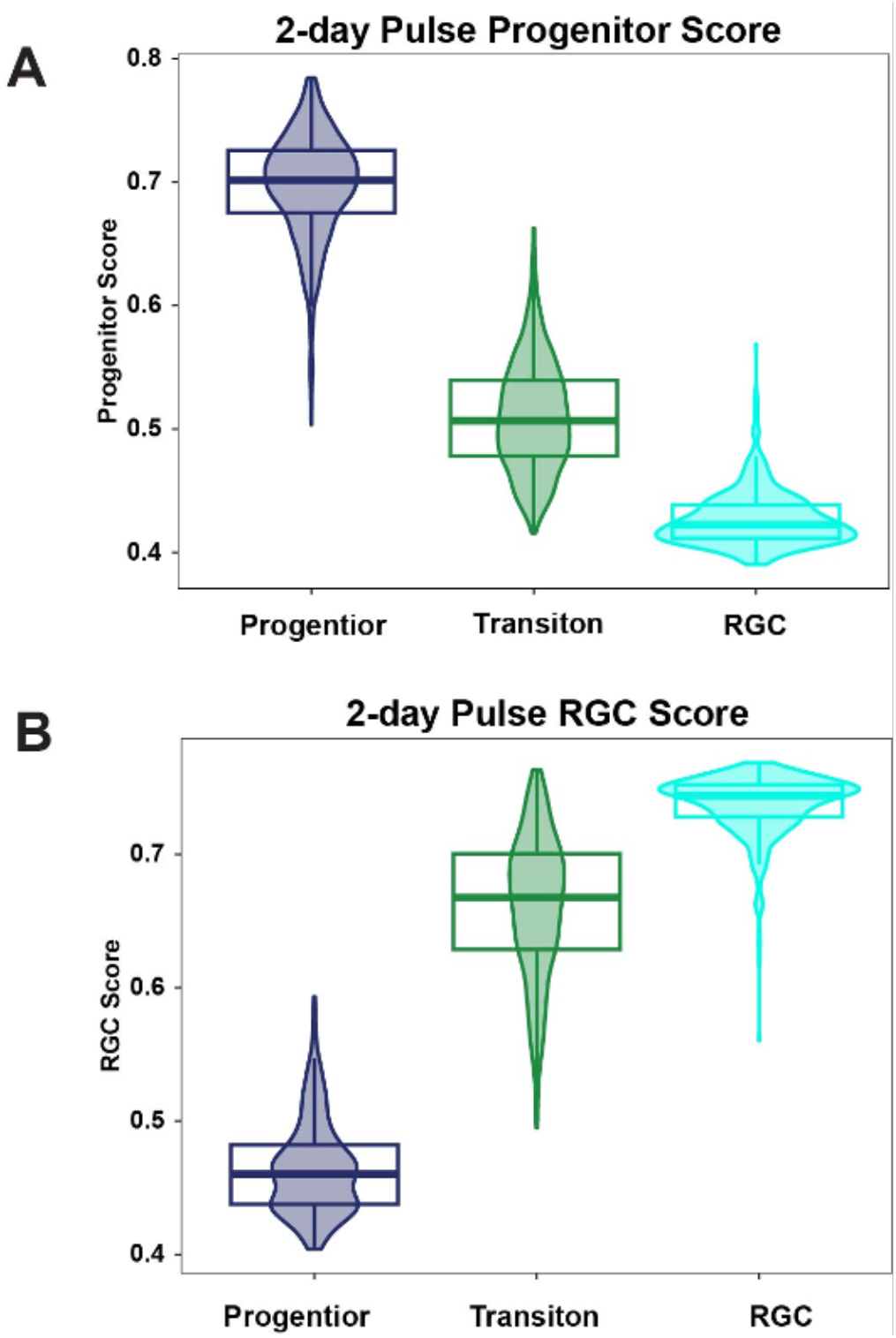
Related to Figure 4. A) Violin plot summarizing the average progenitor gene module score for cells in each cluster of the 2-day pulse shown in **Fig. 4E**. B) Violin plot summarizing the average RGC gene module score for cells in each cluster of the 2-day pulse shown in **Fig. 4E**.

**Supplementary Figure 4.**
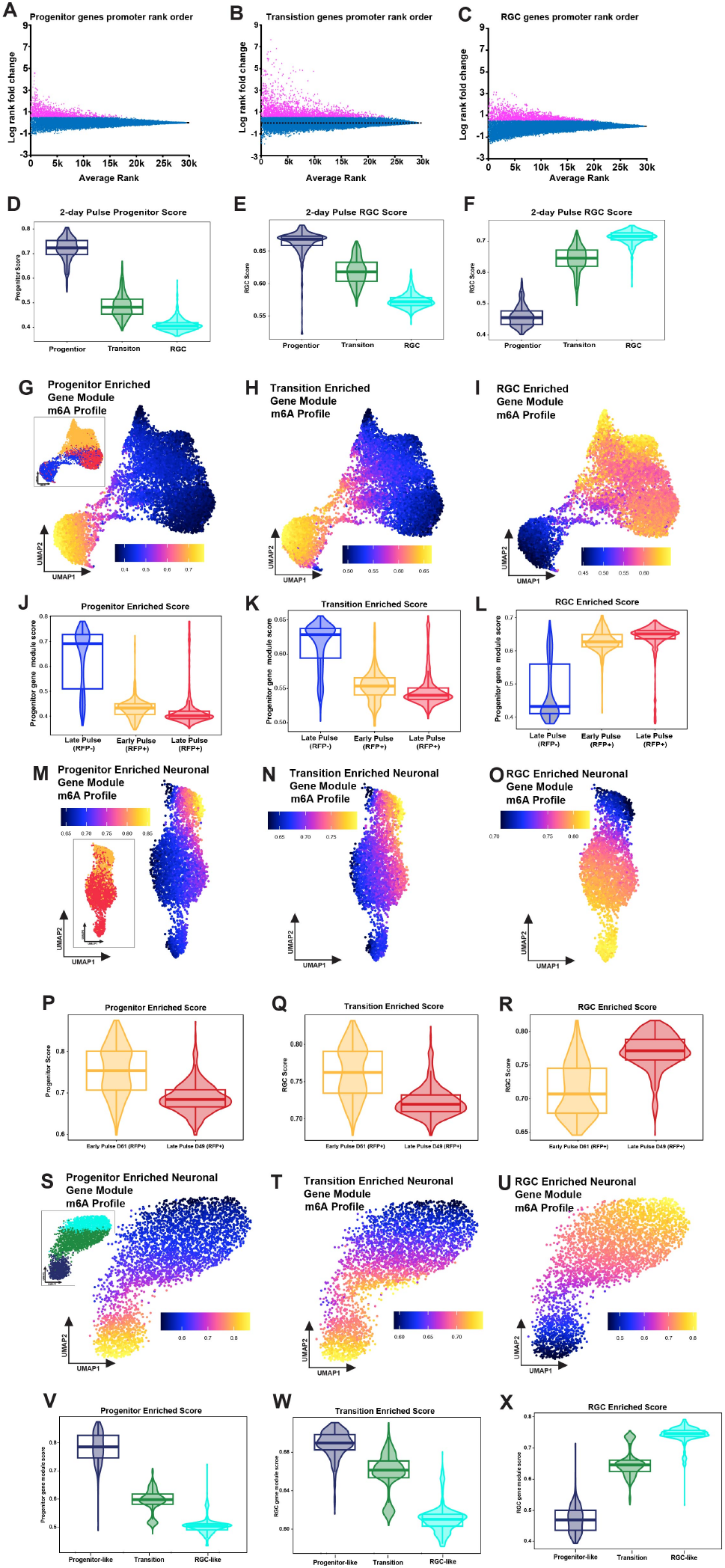
Related to Figure 5. A-C) Promoter rank order of all genes. Pink represents all enriched genes. Blue represents all non-enriched genes as defined in the methods. A) Progenitor gene promotor rank order B) Transition gene promotor rank order C) RGC gene promotor rank order D-F) Violin plots summarizing the average enriched gene module scores for cells in each cluster of the 2-day pulse shown in **Fig. 5A**. D) Progenitor enriched gene module score E) Transition enriched gene module score F) RGC enriched gene module score G-I) UMAP of m6A profiles generated by CUT&TIME for retinal cells cultured with experimental protocol displayed in **Fig. 3A**. G) Gene module of progenitor enriched genes listed in **Table S3**. Inset displays the distribution of FACS sorted cells: Early pulse RFP+ ganglion cells (orange), Late pulse RFP+ ganglion cells (red), and Late pulse RFP-progenitor cells (bule). H) Gene module of transition enriched genes listed in **Table S3**. I) Gene module of RGC enriched genes listed in **Table S3**. J-L) Violin plots summarizing the average enriched gene module scores for cells in each cluster of the 2-day pulse shown in **Fig. S4G-I**. J) Progenitor enriched gene module score K) Transition enriched gene module score L) RGC enriched gene module score M-O) UMAP of m6A profiles generated by CUT&TIME for retinal cells cultured with experimental protocol displayed in **Fig. S2E**. M) Gene module of progenitor enriched genes listed in **Table S3**. Inset displays the distribution of FACS sorted cells: D61 early pulse RFP+ ganglion cells (orange), and D49 late pulse RFP+ cells (red). N) Gene module of transition enriched genes listed in **Table S3**. O) Gene module of RGC enriched genes listed in **Table S3**. P-R) Violin plots summarizing the average enriched gene module scores for cells in the D61 early pulse RFP+ ganglion cells (orange), and D49 late pulse RFP+ cells (red) shown in **Fig. S4M-O**. P) Progenitor enriched gene module score Q) Transition enriched gene module score R) RGC enriched gene module score S-U) UMAP of m6A profiles generated by CUT&TIME for retinal cells cultured with experimental protocol displayed in **Fig. 4A**. S) Gene module of progenitor enriched genes listed in **Table S3**. Inset displays three different clusters; progenitor (Prog, blue), transition (Trans, green), and RGCs (RGC, cyan). T) Gene module of transition enriched genes listed in **Table S3**. U) Gene module of RGC enriched genes listed in **Table S3**. V-W) Violin plots summarizing the average enriched gene module scores shown in **Fig. S4S-O** for retinal cells in the long pulse. V) Progenitor enriched gene module score W) Transition enriched gene module score X) RGC enriched gene module score

